# Methods for the targeted sequencing and analysis of integrons and their gene cassettes from complex microbial communities

**DOI:** 10.1101/2021.09.08.459516

**Authors:** Timothy M. Ghaly, Anahit Penesyan, Alexander Pritchard, Qin Qi, Vaheesan Rajabal, Sasha G. Tetu, Michael R. Gillings

## Abstract

Integrons are bacterial genetic elements that can integrate mobile gene cassettes. They are mostly known for spreading antibiotic resistance cassettes among human pathogens. However, beyond clinical settings, gene cassettes encode an extraordinarily diverse range of functions important for bacterial adaptation. The recovery and sequencing of cassettes has promising applications, including: surveillance of clinically important genes, particularly antibiotic resistance determinants; investigating the functional diversity of integron-carrying bacteria; and novel enzyme discovery. Although gene cassettes can be directly recovered using PCR, there are no standardised methods for their amplification and, importantly, for validating sequences as genuine integron gene cassettes. Here, we present reproducible methods for the PCR amplification, sequence processing, and validation of gene cassette amplicons from complex communities. We describe two different PCR assays that either amplify cassettes together with integron integrases, or gene cassettes together within cassette arrays. We compare the use of Nanopore and Illumina sequencing, and present bioinformatic pipelines that filter sequences to ensure that they represent amplicons from genuine integrons. Using a diverse set of environmental DNAs, we show that our approach can consistently recover thousands of unique cassettes per sample and up to hundreds of different integron integrases. Recovered cassettes confer a wide range of functions, including antibiotic resistance, with as many as 300 resistance cassettes found in a single sample. In particular, we show that class 1 integrons appear to be collecting and concentrating antibiotic resistance genes out of the broader diversity of cassette functions. The methods described here can be applied to any environmental or clinical microbiome sample.

## Introduction

Integrons are bacterial genetic elements that can capture, mobilise and rearrange gene cassettes [1, 2]. They are mostly known for spreading a diverse repertoire of gene cassettes that collectively confer resistance to almost all classes of antibiotics [3]. Beyond clinical settings, however, integrons play a crucial role in bacterial evolution by rapidly generating genomic diversity [4, 5]. Functional integrons are characterised by their flagship gene, the integron integrase (*intI*), which encodes a site-specific tyrosine recombinase (IntI). IntI mediates the insertion of gene cassettes at the integron recombination site (*attI*), which acts as the insertion site of captured gene cassettes [6]. Gene cassettes, prior to their insertion, are circular molecules, which possess a cassette recombination site (*attC*). Their insertion involves IntI-mediated recombination between the *attI* site of the integron and the *attC* site of the cassette [7–11]. Multiple cassettes can be inserted to form a linear cassette array, which can vary in size from zero or one cassette to hundreds [12, 13]. IntI activity is induced by DNA damage, often triggered by environmental stress [14, 15]. Integrons can therefore provide genomic diversity at precisely the moment when it is needed the most, thus facilitating ‘adaptation on demand’ [16].

Recovery and sequence analysis of integron gene cassettes serve several purposes. First, screening gene cassettes can provide a direct method for surveillance of resistance genes that are prevalent in an environment of interest. It has also been proposed that surveying gene cassettes can help detect novel functions that might be harmful to human health, such as increased pathogenicity or resistance to novel antibiotics [17]. In particular, class 1 integrons, due to their mobility, abundance and distribution [18, 19], are primed to play a crucial role in dissemination of these genes. Finally, exploring gene cassettes provides a window into the functional diversity of the bacterial pangenome. Gene cassettes have been found to be extraordinarily abundant and diverse in every environment surveyed [20–26]. Further, many cassettes with known functions act as single-gene/single-trait entities. As such, they need minimal integration into metabolic networks and can likely function in a relatively wide range of genomic contexts. These traits make them highly valuable commodities for synthetic biology and biotechnological applications, particularly for the discovery of diverse enzymatic activities [17].

Currently, gene cassettes can be recovered from genome sequencing of cultured isolates, whole metagenomic sequencing, or amplicon sequencing of *attC*-associated genes. Since most bacteria are yet to be cultured, cassettes identified from isolate genomes inevitably reflect only a small proportion of all gene cassettes, exacerbated by the fact that different strains of the same species can vary widely in cassette content [4]. Whole metagenomic sequencing, although potentially a less biased approach, can be challenging, as gene cassettes often exist at very low abundances and can contain repeat sequences. A targeted amplicon sequencing approach, however, can overcome these issues and could provide the most efficient method for recovering diverse gene cassettes from complex microbial communities [27].

Cassette-targeted amplicon sequencing has been used previously, with varying returns in gene cassette recovery [20–27]. As sequencing technologies have improved, the ability to capture a greater diversity of gene cassettes has also increased [20, 26]. However, such studies lack standardised methods for amplifying and, importantly, validating amplicon sequences as part of genuine cassettes arrays. Here, we present standardised and reproducible methods for amplifying, sequencing, and bioinformatic filtering of genuine gene cassettes from mixed microbial communities.

We applied two different PCR assays using DNA isolated from diverse environmental samples with the aim of recovering integron integrases and gene cassettes. PCR products were sequenced with both long-read Nanopore (ONT) and short-read Illumina MiSeq sequencing technologies. Importantly, we present bioinformatic pipelines that filter sequences for complete *attC* sites or *intI* genes. We show that after filtering, we can consistently recover thousands of gene cassettes from a single sample. We find that recovered genes display a diverse suite of functional traits, including antibiotic resistance.

## Methods

### Sample collection and DNA extraction

Duplicate samples were collected from six different sites (3 x terrestrial and 3 x aquatic environments). Terrestrial sites consisted of urban parkland soil from Macquarie University (Sydney, New South Wales, Australia) [20], hot desert soil from Sturt National Park (North-western New South Wales) [28, 29], and Antarctic soil from Herring Island [20]. The aquatic sites consisted of river sediment (Lane Cove River, New South Wales), freshwater biofilms (Mars Creek, New South Wales) [30], and estuarine sediment (Paramatta River Estuary, New South Wales). From each of the 12 samples, DNA was extracted from 0.3 g of material using a standard bead-beating protocol [31]. Each resulting DNA sample was used as the template for two different PCR assays, described below, and all were subsequently sequenced using long-read Nanopore (ONT) and short-read MiSeq technologies (Fig. 1).

**Figure 1.**
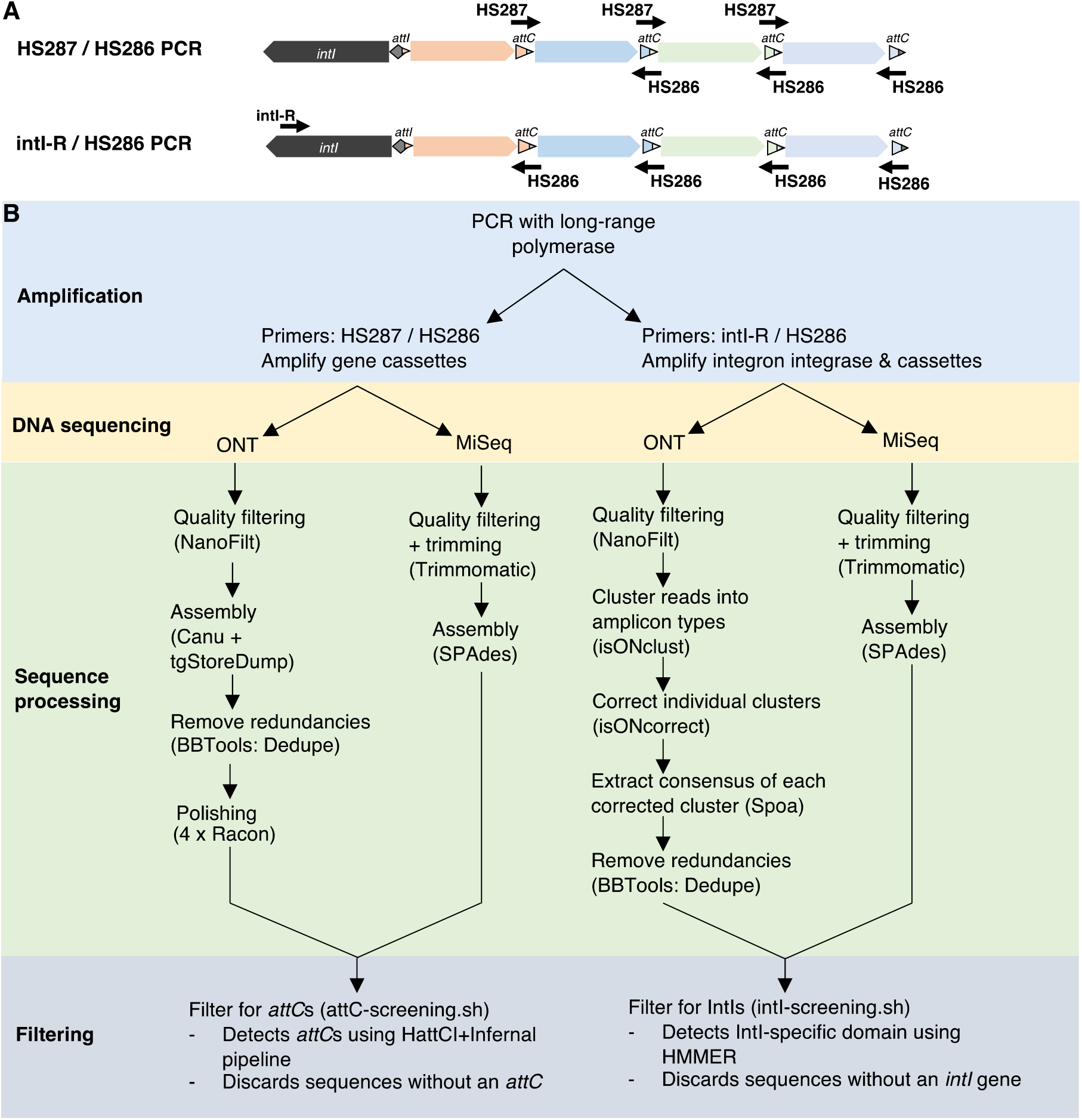
Experimental and bioinformatic workflow for gene cassette amplicon sequencing. (**A**) Components of integrons amplified by the two PCR assays. The primer set HS287 / HS286 targets cassettes that lie between two *attC* sites. Potentially any gene cassette(s) can be amplified by this primer set. The primer set intI-R / HS286 targets diverse integron integrases (*intI*) and cassette recombination sites (*attC*). The resulting amplicons include ~800 bp of *intI* and at least the first cassette(s) of an array. (**B**) The bioinformatic steps and software (in parentheses) used to process and filter amplicon data. Methods are shown for both primer sets sequenced with either Nanopore (ONT) or Illumina MiSeq.

### PCR amplification and DNA sequencing

For each sample, we conducted two different PCR assays (Fig. 1A). The first used the primers HS287 and HS286 [27], which target *attC* recombination sites in opposing directions to amplify intervening gene cassettes. The second primer set, intI-R / HS286, amplifies approximately 800bp of the integron integrase gene as well as adjacent gene cassettes. The primer intI-R (5’-GCG AAC GAR TGB CGV AGV GTG TG −3’) was designed to target diverse integron integrases and was based on an alignment of 174 complete *intI* sequences containing a functional catalytic site, as compiled by Cambray et al. [14]. Importantly, the last 6 bp of the 3’ end of intI-R exactly matches 75% of aligned *intI* sequences. For amplification, we used Phusion Hot Start II DNA Polymerase (ThermoFisher Scientific, Waltham, MA, USA), which is a long-range DNA polymerase, chosen to facilitate the amplification of large segments of integron cassette arrays, known to reach more than 100 kilobases in length [12]. The PCRs were carried out in 50 μL volumes containing a final concentration of 1 x GC Phusion Buffer, 0.2 mM dNTPs, 0.5 μM of each primer, 3% DMSO and 2 U of Phusion DNA polymerase. All PCRs were performed using GeneReleaser® (Bioventures, Murfreesboro, TN, USA) as previously described [32]. Triplicate PCRs were performed for each sample to increase the chances of capturing rare gene cassettes that might otherwise escape amplification due to the stochastic nature of PCR.

For the HS287 / HS286 primer set, the following thermal cycling program was used: 98°C for 3 min for 1 cycle; 98°C for 10 s, 60°C for 30 s, 72°C for 3 min 30 s for 35 cycles; and a final extension step at 72°C for 10 min. For the intI-R / HS286 primer set, the following thermal cycling program was used: 98°C for 3 min for 1 cycle; 98°C for 10 s, 65°C for 30 s, 72°C for 3 min 30 s for 35 cycles; and a final extension step at 72°C for 10 min. PCR efficiency was assessed using 2% agarose gel electrophoresis. Triplicate PCRs were pooled and then purified with AMPure XP beads (Beckman Coulter, Danvers, MA, USA) according to the manufacturer’s protocol.

For long-read sequencing, the 24 PCR products (representing the 12 samples amplified with each primer set) were multiplexed in a single DNA library using the ONT Ligation Sequencing Kit (SQK-LSK109) and the ONT Native Barcoding Expansion Kits (EXP-NBD104 and EXP-NBD114) according to the manufacturer’s protocol. The DNA library was sequenced using a MinION MK 1B sequencing device on an R10.3 flow cell. Sequencing was allowed to run for 24 hours. Basecalling was carried out with Guppy v.4.3.4 with default parameters using the high accuracy (HAC) basecalling model.

For short-read sequencing, the 24 PCR products underwent an Illumina DNA shotgun library preparation using the Nextera XT protocol and then sequenced with MiSeq 300 bp paired-end sequencing on a single lane. Illumina sequencing and library preparation were carried out at the Australian Genome Research Facility (Melbourne, Australia).

### Sequence processing and attC filtering: HS287 / HS286 PCRs

To compare short- and long-read sequencing technologies, we sequenced HS287 / HS286 PCRs on both Nanopore (ONT) and MiSeq platforms. The respective workflows and software used for sequence processing and filtering are summarised in Figure 1B.

For ONT sequences of the HS287 / HS286 PCRs, we first filtered reads based on average quality (q) scores. We removed any reads with an average q score below 10 using NanoFilt v2.8 [33] [parameters: -q 10]. Although each read spans the length of an entire amplicon, many amplicons represent overlapping subsections of larger potential templates. Thus, an assembly of these initial reads into larger cassette arrays was carried out using Canu v2.0 [34] [parameters: genomeSize=5m minReadLength=250 minOverlapLength=200 corMinCoverage=0 corOutCoverage=20000 corMhapSensitivity=high maxInputCoverage=20000 batMemory=125 redMemory=32 oeaMemory=32 batThreads=24 purgeOverlaps=aggressive]. Assembled contigs and unassembled reads were then extracted together using the tgStoreDump script within Canu [parameters: -consensus -fasta]. Any redundancies were removed using dedupe.sh, available from the BBTools package v35 [35] with default parameters. Consensus sequences were then corrected with 4 rounds of polishing using Racon v1.4.20 [36]. Each round of Racon polishing involved read mapping with minimap2 v2.20-r1061 [37] [parameters: -x map-ont -t 24] and error correction with Racon [parameters: -m 8 -x 6 -g -8 -w 500 -t 24].

For MiSeq sequence data of the HS287 / HS286 PCRs, paired-end reads first underwent quality trimming and adapter clipping using Trimmomatic v0.38 [38] [parameters: -phred33 ILLUMINACLIP:adapters.fa:2:30:10 LEADING:3 TRAILING:3 SLIDINGWINDOW:4:15 MINLEN:30] where ‘adapters.fa’ is a fasta-formatted file containing all commonly used Illumina adapter sequences. If only one end of paired reads had acceptable quality, it was used as a single read during assembly. Surviving paired-end reads and single reads were assembled together using SPAdes v3.14.1 [39–41] [parameters -k 21,33,55,77,99,127 --only-assembler --careful].

The resulting ONT and MiSeq sequences were both filtered based on the presence of internal cassette recombination sites (*attC*s). Given the degenerate nature of the primers, off-target amplicons might constitute a significant portion of the reads. Filtering for sequences that have internal *attC* sites is thus an essential step when analysing cassette amplicon data. It should be noted that filtering for *attC*s in this way may discard some genuine amplicons which consist of single gene cassettes, since they do not possess a complete *attC* site (Fig. 1A). Nevertheless, we consider that for obtaining meaningful ecological data, the removal of potential false positives is more important than the loss of some true positives. To filter for *attC* sites, we used an in-house script, attC-screening.sh (available: https://github.com/timghaly/integron-filtering), with default parameters. The script uses the HattCI [42] + Infernal [43] pipeline that has been previously described [44, 45]. In short, attC-screening.sh searches for the sequence and secondary structures conserved among *attC*s and retains any input sequence that has at least one *attC* site. The script can be used on data generated from any sequencing technology.

### intI-R / HS286 PCRs: sequence processing and IntI filtering

All intI-R / HS286 PCRs were also sequenced on both ONT and MiSeq platforms (Fig. 1B). For Nanopore sequencing, basecalled reads were first quality filtered using NanoFilt v2.8 [33] [parameters: -q 10]. Reads representing concatemers and chimeras were removed using yacrd v0.6.2 [46] with default parameters for ONT data. Since all amplicons should be anchored on one end to the *intI* gene, an assembly would not be suitable. Instead, we clustered reads into amplicon ‘types’ using isONclust v0.0.6.1 [47]. Each cluster was then individually corrected using isONcorrect v0.0.8 [48] with default parameters. Unlike error-correction of genomic data, isONcorrect takes into account uneven coverage within the same read as well as structural variation among similar reads from different clusters (e.g., reads that represent true biological rearrangements of the same gene cassettes). From each corrected cluster, a consensus sequence was then generated using spoa v4.0.7 [36] [parameter: -r 0]. Any redundancies were removed using the BBTools v35 [35] script dedupe.sh with default parameters.

MiSeq sequences were processed in the same manner as described above for the HS287 / HS286 data. This involved quality trimming and adapter clipping using Trimmomatic v0.38 [38], followed by an assembly of the reads using SPAdes v3.14.1 [39–41].

The resulting ONT and MiSeq sequences were both filtered based on the presence of IntI protein sequences. To detect sequences that encoded IntI, we used an in-house script, intI-screening.sh (available: https://github.com/timghaly/integron-filtering), with default parameters. The script uses a profile hidden Markov model (HMM) provided by Cury et al. [13] to detect the additional domain that is unique to integron integrases, separating them from other tyrosine recombinases [49]. The intI-screening.sh pipeline first uses Prodigal [50] to predict all encoded protein sequences, and then screens them for the IntI-specific domain using hmmsearch from the HMMER v3 software package [51]. Any sequences that do not contain a recognisable integron integrase are discarded. Similarly, intI-screening.sh can be used on data generated from any sequencing technology.

### Protein prediction and functional classification of gene cassettes

Cassette open reading frames (ORFs) and their translated protein sequences were predicted using Prodigal v2.6.3 [50] in metagenomic mode [parameters: -p meta].

To assess the broad-scale functional diversity of gene cassettes, we used the Clusters of Orthologs Groups (COGs) database [52]. COG functions were assigned to cassette-encoded protein sequences using eggNOG-mapper v2.0.1b [53, 54] executed in DIAMOND [55] mode with default parameters. To detect cassette-encoded antimicrobial resistance genes (ARGs), we used ABRicate v0.8 [56] to search against the Comprehensive Antibiotic Resistance Database (CARD) [57] [Downloaded: 2021-Apr-21].

### Taxonomic classification of attC sites

The gene cassettes of sedentary chromosomal integrons (SCIs) generally possess highly similar *attC* sites, and this conservation spans the SCIs of different bacteria within the same taxon [4, 58, 59]. We have recently modelled the conserved sequence and structure of *attC* sites from the chromosomal integrons of 11 bacterial taxa [44]. These included six Gammaproteobacterial orders (Alteromonadales, Methylococcales, Oceanospirillales, Pseudomonadales, Vibrionales, Xanthomonadales) and an additional five phyla (Acidobacteria, Cyanobacteria, Deltaproteobacteria, Planctomycetes, Spirochaetes). A covariance model (CM) was generated separately for each taxon, and this can be used to correctly identify the source taxon of *attC* sites with high specificity (98-100%) [44].

Here, we used an in-house script, attC-taxa.sh (available: https://github.com/timghaly/attC-taxa), with default parameters to detect any *attC* sites that have originated in the SCIs of one of the 11 taxa. The attC-taxa.sh pipeline incorporates all 11 CMs and uses cmsearch [parameters: --notrunc --max] from the Infernal software package [43] to classify *attC*s. It is important to note that each taxon-specific model exhibits different sensitivities in detecting true positives and thus the relative proportion of different taxa cannot be compared within the same sample. However, the relative proportion of the same taxon can be compared between different samples.

### ONT – MiSeq comparisons

For comparisons of the cassette and integrase diversity recovered between ONT and MiSeq technologies, we first considered differences in sequencing depth. To do this, we randomly subsampled 50 Mb of raw reads from each sample using rasusa v0.3.0 [60] [parameters: -- coverage 50 --genome-size 1Mb]. After subsampling, all sequence processing and filtering steps were repeated as described above.

All formal comparisons were made using two-sample T-tests (or Wilcoxon rank sum tests if the data were not normally distributed) using the rstatix v0.7.0 R package [61]. To determine if the data were normally distributed, Shapiro-Wilk tests were carried out using rstatix v0.7.0 [61], as well as visually inspected the distributions against their theoretical normal distributions using Q-Q plots generated with the R package ggpubr v0.4.0 [62].

To assess the overlap in recovered ORFs between ONT and MiSeq, we mapped the cassette ORFs from one technology to the reads of the other using minimap2 v2.20-r1061 [37]. We considered the ORF to be present if it had a mean coverage depth of at least 1x that spanned at least 98% of the ORF. For ONT and MiSeq read mapping, we used the minimap2 presets [-ax map-ont -t 8] and [-ax sr -t 8], respectively. Coverage statistics were extracted from the mapping alignments using the ‘sort’ and ‘coverage’ programs within the SAMtools software package [63, 64].

## Results and Discussion

Here, we present a stringent pipeline for PCR amplifying, sequencing and analysing integron integrases and gene cassettes from diverse microbial communities (Fig. 1). For this, we used two different PCR primer sets, HS287 / HS286 and intI-R / HS286 (Fig. 1A). The sample types consisted of a wide variety of soils (from an urban parkland, an Australian desert, and an Antarctic island), as well as river and estuarine sediments, and freshwater biofilms.

To assess the suitability of long- and short-read sequencing technologies, we sequenced amplicons from both PCR assays using Nanopore (ONT) and Illumina MiSeq platforms, respectively. Average ONT yield was 181 Mb (100 – 358 Mb per sample) for the HS287 / HS286 primer set and 216 Mb (62 – 502 Mb per sample) for the intI-R / HS286 primer set. The average MiSeq yield was 418 Mb (228 – 720 Mb per sample) and 663 Mb (275 – 1,247 Mb per sample), respectively for these primer sets.

To ensure amplicons were part of genuine integrons, we filtered the HS287 / HS286 data for *attC* sites, and the intI-R / HS286 data for IntI protein sequences (Fig. 1B). For the HS287 / HS286 data, an average of 23.8% and 19.0% of amplicon sequences were retained after filtering for ONT and MiSeq, respectively (Supplementary Fig. S1A). While, for the intI-R / HS286 data, an average of 1.2% and 1.5% of sequences remained after filtering for ONT and MiSeq, respectively (Supplementary Fig. S1B). The difference in proportions of sequences retained after filtering between ONT and MiSeq were not statistically significant for either primer set. The low proportion of surviving sequences for the intI-R / HS286 data is likely a result of the intI-R primer binding to other tyrosine recombinases. While many sequences were filtered out, the data retained from this primer set, as described below, include a large, diverse set of both known and entirely novel integron integrases and gene cassettes.

The lengths of the recovered sequences for both primer sets were significantly larger for ONT sequencing compared to MiSeq (Supplementary Fig. S2). For the HS287 / HS286 set, sequence lengths ranged from 500 bp to more than 25,304 bp for ONT, and 500 bp to 19,244 bp for MiSeq. For the intI-R / HS286 data, sequence lengths ranged from 803bp to 16,179bp for ONT and 800bp to 7,432bp for MiSeq.

### Recovered diversity of gene cassette ORFs

We assessed the efficiency of both primer sets in recovering gene cassette open reading frames (ORFs). Among all 12 samples, the HS287 / HS286 primers amplified 33,854 and 62,118 non-redundant cassette-encoded proteins when sequenced with ONT and MiSeq, respectively (Fig. 2A). After adjusting for sequencing depth, there was no significant difference in cassette recovery between the two sequencing technologies (Fig. 2B). On average, we observed that ~50% of cassette ORFs sequenced with one technology were also recovered by the other (Supplementary Fig. S3A).

**Figure 2.**
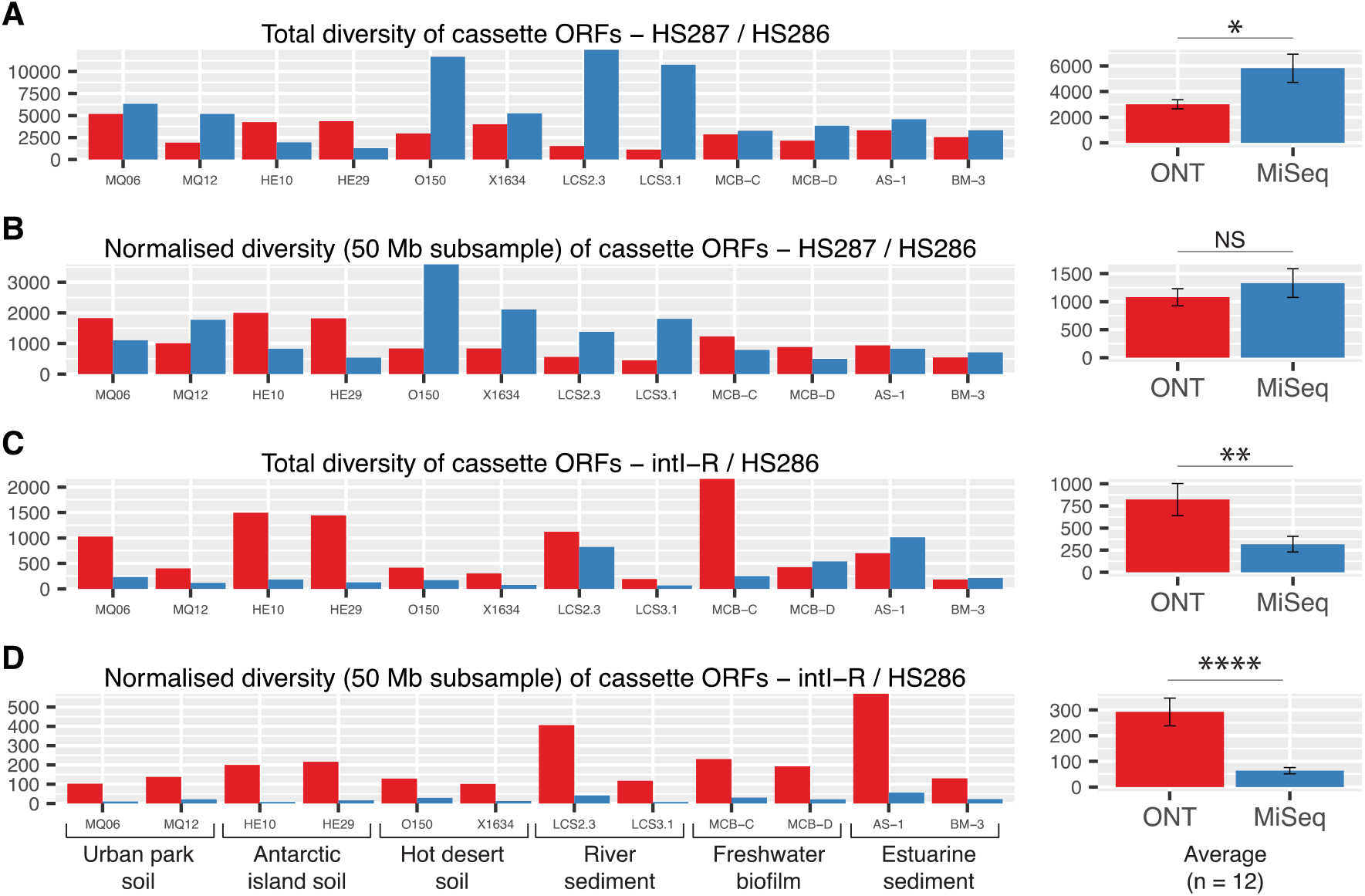
Diversity of recovered gene cassette ORFs. Redundancy was removed using a 100% amino acid identity of translated protein sequences. (**A**) Total non-redundant cassette ORFs amplified using the primers HS287 / HS286. (**B**) Cassette ORF diversity was normalised for HS287 / HS286 sequencing depth (based on a 50 Mb subsample of sequence reads). Total (**C**) and normalised (**D**) cassette ORF diversity are shown for the intI-R / HS286 primer set. Average (± 1 S.E) diversity for each analysis are shown on the right-hand side of each panel. The degree of statistical significance is shown by asterisks as determined by two-sample T-tests or Wilcoxon rank sum tests (depending on the normality of the data). NS: P>0.05, *: P <0.05, **: P<0.01, ***: P<0.001, ****: P<0.0001.

The HS287 / HS286 primer set is preferred in order to recover the greatest diversity of gene cassettes. Indeed, the recovery rate of gene cassettes using the methods described here surpasses any previously described approach. Notably, Pereira et al. [45] conducted an impressive survey of gene cassettes from 10 terabases of metagenomic data obtained from 14 public databases. Across all datasets, they identified an average of 0.03 unique cassette ORFs per 500 kilobases of assembled data. Here, we recover 218 and 265 ORFs per 500 kilobases of assembled data when sequenced with ONT and MiSeq, respectively. Although screening metagenomes may provide a relatively unbiased approach in analysing gene cassettes, it clearly requires much deeper sequencing to recover sufficient cassette data for in-depth ecological or evolutionary analyses. Studies of integrons and their associated genetic cargo will therefore continue to benefit from the use of amplicon sequencing approaches, such as described in the present study.

For the intI-R / HS286 primer set, we recovered a total of 9,641 and 3,742 non-redundant cassette ORFs when sequenced with ONT and MiSeq, respectively (Fig. 2C). ONT sequencing recovered significantly more (P<0.0001) of the cassette ORF diversity of each sample than MiSeq (Fig. 2D). This was despite all ONT cassette ORFs being covered by the MiSeq reads (Supplementary Fig. S3B). This shows that although the MiSeq reads cover all the cassettes being amplified by the primers, recovery of the cassettes is sub-optimal, most likely due to difficulties in their assembly. In particular, different cassette arrays associated with the same or similar integron integrase are likely to be problematic for a short-read assembly approach. While the intI-R / HS286 primer pair does not recover as much diversity as the HS287 / HS286 set, it does provide additional key information on IntI diversity (discussed further below) and indicates which gene cassettes are associated with which *intI* genes.

The intI-R / HS286 primer set can also reveal which gene cassettes are located towards the start of a cassette array (Fig. 1A). This is of biological and ecological significance, since the first cassettes in arrays are the most recently inserted cassettes and are likely to be strongly expressed [65]. During environmental perturbations, integron integrase activity leads to the acquisition of novel and rearrangement of existing cassettes, inserting them at the start of the array where strong expression is guaranteed [17, 66, 67]. Selection fixes lineages with first-position cassettes that confer significant advantages. Thus, gene cassettes recovered from the intI-R / HS286 primer set might provide important ecological insights at the time of sampling.

### Recovered diversity of integron integrases

Using the intI-R / HS286 primers, we recovered a total of 1,413 and 1,867 different integron integrase genes among the 12 samples when sequenced with ONT and MiSeq, respectively (Fig. 3A). There was no significant difference in integron-integrase recovery between the two sequencing technologies, with or without adjusting for sequencing depth (Figs. 3A-B). Both sequencing technologies could recover an impressive diversity of IntIs from the 12 samples. In comparison, a comprehensive screening of 2,484 bacterial genomes recovered only 215 different IntIs [13]. This shows that despite the low rate of intI-R / HS286 sequences that are retained after filtering, a significant number of novel integron sequences are recovered.

**Figure 3.**
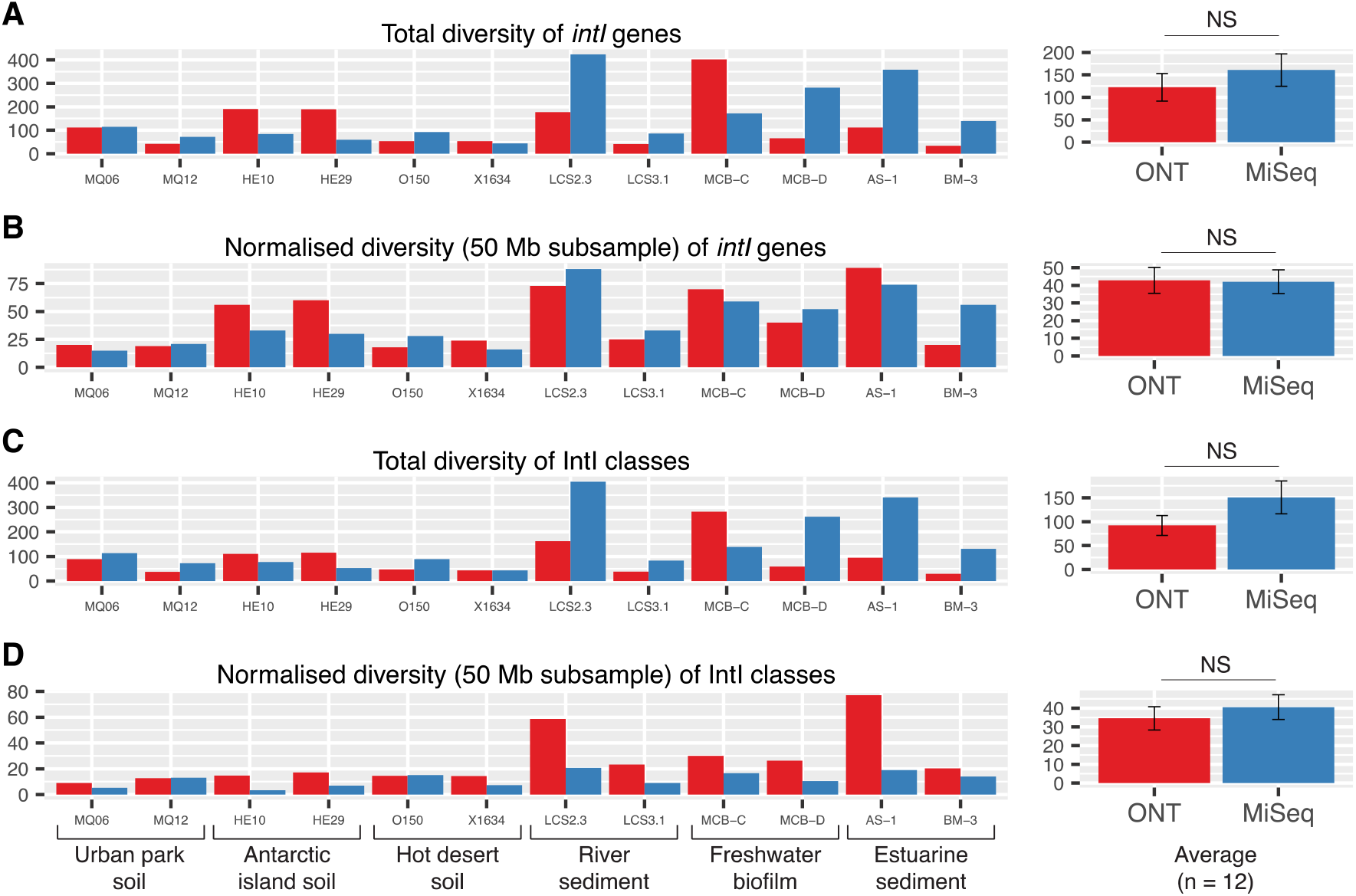
Diversity of integron integrases recovered by the intI-R / HS286 primer set. (**A**) Total non-redundant (100% amino acid identity) integron integrases (IntIs) recovered. (**B**) IntI diversity was normalised for sequencing depth (based on a 50 Mb subsample of sequence reads). Total (**C**) and normalised (**D**) diversity of IntI classes (using a 94% amino acid clustering threshold) are shown. Average (± 1 S.E) diversity for each analysis are shown on the right-hand side of each panel. Differences between Nanopore (ONT) and Illumina MiSeq technologies were not significant (NS) as determined by Wilcoxon rank sum tests.

To determine how many classes of integrons these IntIs represented, we sought to define the amino acid clustering threshold for an integron class. To do this, we used the most abundant and widely distributed IntI, the class 1 integron integrase (IntI1) [68]. Here, we iteratively set decreasing amino acid clustering thresholds for our library of IntIs using CD-HIT v4.6 [69, 70] [parameters: -n 5 -d 0 -g 1 -t 0]. We continued until all IntI1s in our dataset were grouped into a single cluster while ensuring all non-IntI1s were excluded (Supplementary Fig. S4A). This resulted in a 94% amino acid identity being selected as the most appropriate clustering threshold for IntI1s. Although this might not reflect the ideal threshold for all classes, it nevertheless provides a semi-quantitative approach to defining an integron class based on amino acid homology.

Using a 94% clustering threshold, we recovered a total of 984 and 1,646 integron classes among our dataset when sequenced with ONT and MiSeq, respectively (Supplementary Fig. S4B). There was no significant difference in integron class recovery between the two sequencing technologies, with or without adjusting for sequencing depth (Figs. 3C-D). In addition, we examined the most prevalent integron classes; defined here as those being IntIs that were present in at least one-third of all samples (Table 1). This identified ten prevalent IntI classes, found to be 60-70% similar to endogenous IntIs from diverse bacterial phyla (Table 1). Not surprisingly, class 1 integrons were the only class to be found in every sample, including those from Antarctica and outback Australia.

**Table 1.** Most prevalent integron integrase (IntI) classes.

| Number of IntIs in cluster/class | Percentage of samples present in | BLASTP taxa | BLASTP amino acid % |
| --- | --- | --- | --- |
| 94 | 100 | Class 1 integron - Multispecies | 99.3 |
| 71 | 91.7 | Xanthomonadales<br>(Rhodanobacteraceae & Xanthomonadaceae) | ~70 |
| 19 | 91.7 | Multiple phyla (Deltaproteobacteria, Nitrospinae, Chloroflexi) | ~60 |
| 26 | 66.7 | Xanthomonadaceae (*Lysobacter*, *Vulcaniibacterium*, *Thermomonas*, *Luteimonas*, *Pseudoxanthomonas*) | ~70 |
| 10 | 58.3 | Betaproteobacteria | ~80 |
| 13 | 41.7 | 'IntI1-like' - Multispecies | ~91 |
| 13 | 41.7 | Rhodanobacteraceae | ~75 |
| 12 | 41.7 | Xanthomonadales<br>(Rhodanobacteraceae & Xanthomonadaceae) | ~74 |
| 8 | 41.7 | Xanthomonadales<br>(Rhodanobacteraceae & Xanthomonadaceae) | ~72 |
| 7 | 41.7 | Planctomycetes | ~67 |

### Functional diversity of gene cassettes

Here we show that gene cassette ORFs largely encode proteins of unknown functions (Fig. 4A). This is in agreement with previous functional analyses of gene cassettes [5, 20, 24, 25]. On average, only ~20% of cassette-encoded proteins amplified with HS287 / HS286 could be assigned a COG functional category, approximately half of which could be assigned a category of known function (Fig. 4A). The dominant COG categories were ‘Transcription’, ‘Replication, recombination and repair’, and ‘Amino acid transport and metabolism’ (Fig. 4B). We show that our methods are capable of recovering gene cassettes that confer a wide range of traits spanning many functional classes.

**Figure 4.**
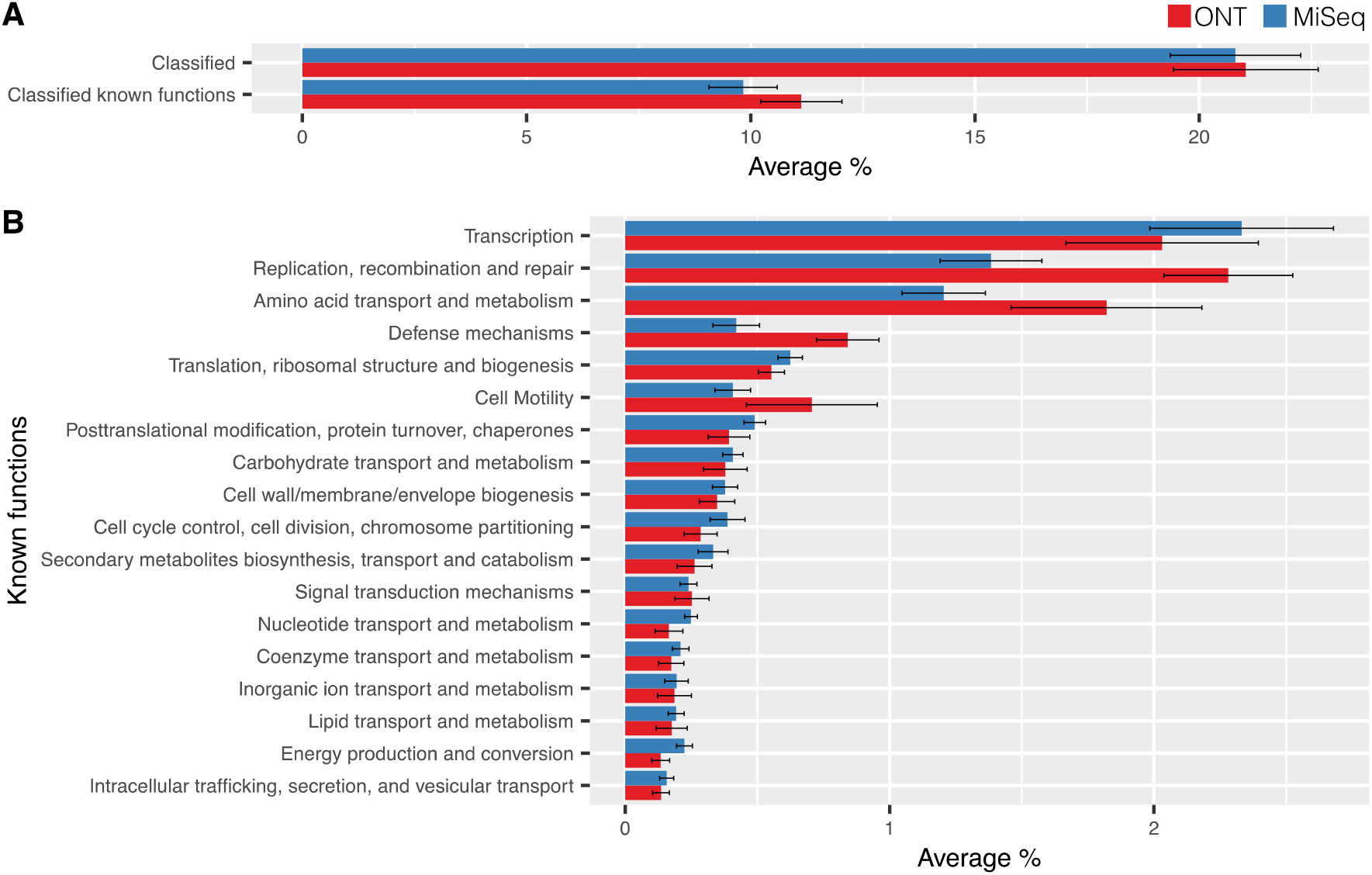
COG functional analysis of cassette-encoded proteins recovered with the HS287 / HS286 primer set. (**A**) Average (± 1 S.E) percentage of proteins per sample (n=12) that can be classified into functional categories. On average ~20% of cassette-encoded proteins can be classified by a COG category, half of which fall into categories of known function. (**B**) The average (± 1 S.E) proportion of proteins within a sample assigned to each of the known functional categories.

### Cassette-encoded antibiotic resistance

For a more specific functional characterisation, we focused on antimicrobial resistance, since these phenotypes are often conferred by integron gene cassettes in clinical settings [3, 71, 72]. Interestingly, we found that for either primer set, ONT sequencing could recover many more ARGs than MiSeq, the latter recovering no ARGs for most samples (Fig. 5A). In contrast, ONT sequencing recovered as many as 300 ARG cassettes within a single sample. We suspect that this discrepancy is an artefact caused by the high similarity between different ARG types, and multiple arrangements of the same ARGs in class 1 cassette arrays that makes their assembly difficult from short-read data. In total, we recovered 106 different ARGs from both primer sets when sequenced with ONT (Fig. 5B). Almost all ARG cassettes encoded proteins known to confer resistance to β-lactam and aminoglycoside antibiotics, these being the most commonly observed integron-mediated resistance types [3].

**Figure 5.**
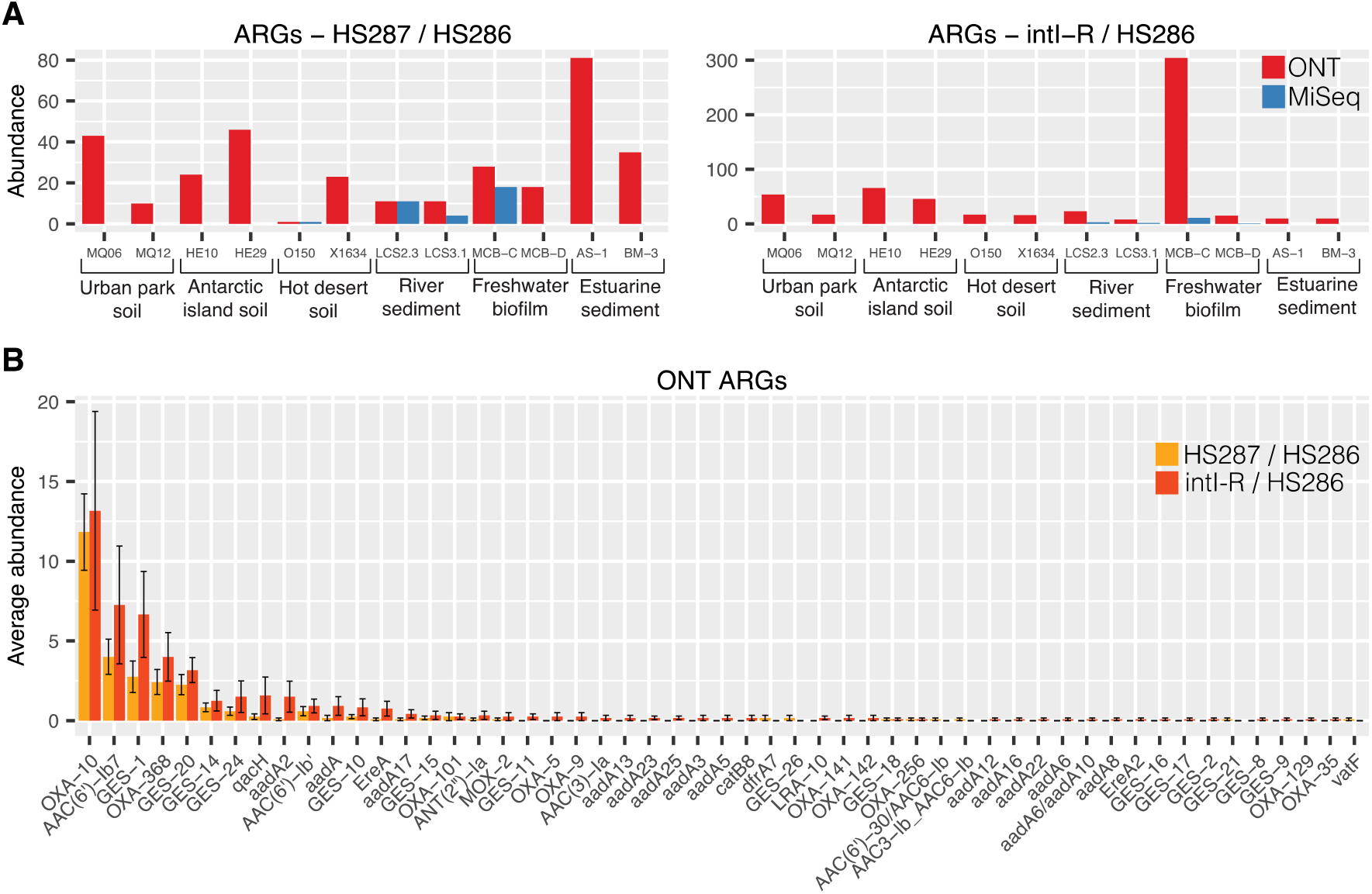
Abundance and diversity of antibiotic resistance gene (ARG) cassettes. (**A**) Abundance of ARGs recovered from either primer set. (B) The average (± 1 S.E) abundance of each ARG type recovered from Nanopore (ONT) sequencing per sample (n=12).

Upon examining all cassette ORFs associated with class 1 integron integrases recovered using intI-R / HS286 primers (Table 1), we found that 162 of 462 (34.6%) were known ARGs. In comparison, only 586 of the 10,385 (5.6%) total cassette ORFs amplified with this primer set were known ARGs. These findings show that class 1 integrons are collecting and concentrating ARG cassettes out of the broader diversity of cassette functions. This enrichment strongly supports the idea that class 1 integrons are key vectors for acquisition and dissemination of antibiotic resistance [3, 73, 74].

### Taxonomic classification of attC sites

We could identify the likely taxonomic sources of 5,998 *attC*s (18.8%) and 10,257 *attC*s (20%) sequenced with ONT and MiSeq, respectively. For taxonomic classification, we used models that capture the sequence and structural homology of chromosomal *attC*s from 11 different taxa. These included six Gammaproteobacterial orders (Alteromonadales, Methylococcales, Oceanospirillales, Pseudomonadales, Vibrionales, Xanthomonadales) and an additional five phyla (Acidobacteria, Cyanobacteria, Deltaproteobacteria, Planctomycetes, Spirochaetes). It should be noted that although the specificity (ability to reject false positives) of each model is very high (98-100%), they exhibit a wide range of sensitivities (proportion of true positive detected) [44]. Therefore, relative abundance of each taxon cannot be compared within the same sample, however, the same taxon can be compared between different samples. It also indicates that the relative abundances of each taxon are likely to be lower-bound estimates.

Here, we show that the relative abundance of each taxon varied across the different sampled environments (Fig. 6). For example, both the Cyanobacteria- and Methylococcales-type *attC*s were most abundant in urban parkland soil (Fig. 6C-D), while Vibrionales-type *attC*s were more abundant in estuarine sediments and freshwater biofilm samples (Fig. 6F and Supplementary Figure S5 for a comparison of all 11 taxa). Such data can provide useful information on the taxonomic contribution to gene cassette pools among different samples.

**Figure 6.**
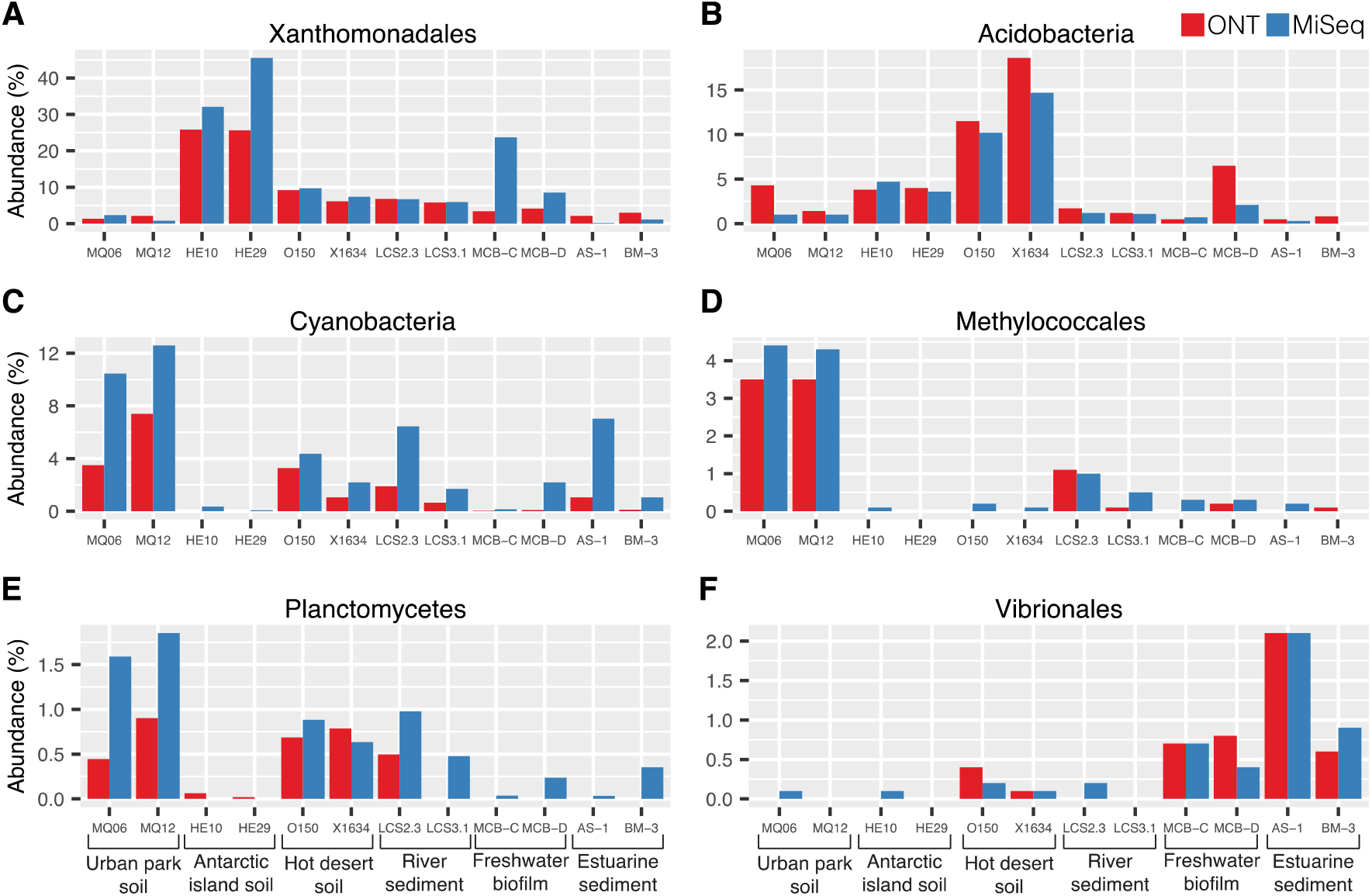
Taxonomic classification of gene cassette recombination sites (*attC*s). Taxonomic predictions are based on a selection of six (**A-F**) of the eleven available taxonomic models of chromosomal *attC*s. Each figure panel shows the proportion of *attC*s across each sample that exhibit sequence and structure conserved among that taxon. For a comparison of all eleven taxa, see Supplementary Figure S5.

## Conclusions

Here, we present experimental and bioinformatic methods for the PCR amplification, DNA sequencing and analysis of integrons from microbial communities. We describe approaches using two different PCR assays and compare the outputs from ONT and MiSeq sequencing. We find that, relative to sequencing depth, ONT generally outperforms or performs the same as MiSeq regarding the recovery of gene cassettes and integron integrases. Most notably, ONT outperforms MiSeq in the recovery of complete ARG gene cassette sequences. We also find that the primer set HS287 / HS286 is efficient at amplifying a wide range of gene cassettes, encompassing extensive *attC* and functional diversity. However, the intI-R / HS286 primer set can provide additional useful information in linking gene cassettes with an integron class. For example, we show that class 1 integrons are collecting and concentrating ARGs relative to the broader cassette pool.

Our described methods can recover key information on the diverse pool of gene cassettes that are helping drive adaptation and niche specialisation in bacteria [4, 16, 67]. Such an approach allows us to investigate the potential traits that are available to integron-carrying bacteria, and to understand the role that gene cassettes play in mediating evolutionary responses under environmental or clinical selection pressures. In addition, the large proportion of cassettes with unknown functions provides an important resource for the discovery of novel enzymatic activities [17].

## Data availability

All sequence data generated in this study are available from the NCBI SRA database under the BioSample accessions SAMN21354384 to SAMN21354431. All BioSamples are linked to the NCBI BioProject PRJNA761546.

## Code availability

The code used for filtering sequences to ensure that they represent amplicons from genuine integrons are available at https://github.com/timghaly/integron-filtering. The code used to predict the taxonomic sources of gene cassette recombination sites (*attC*s) is available at https://github.com/timghaly/attC-taxa.

## Acknowledgements

This research was supported by the Australian Research Council Discovery Grant DP200101874. TMG would like to thank Mary and Saoirse Ghaly for their loving support.

## Supplementary Figures

**Figure S1.**
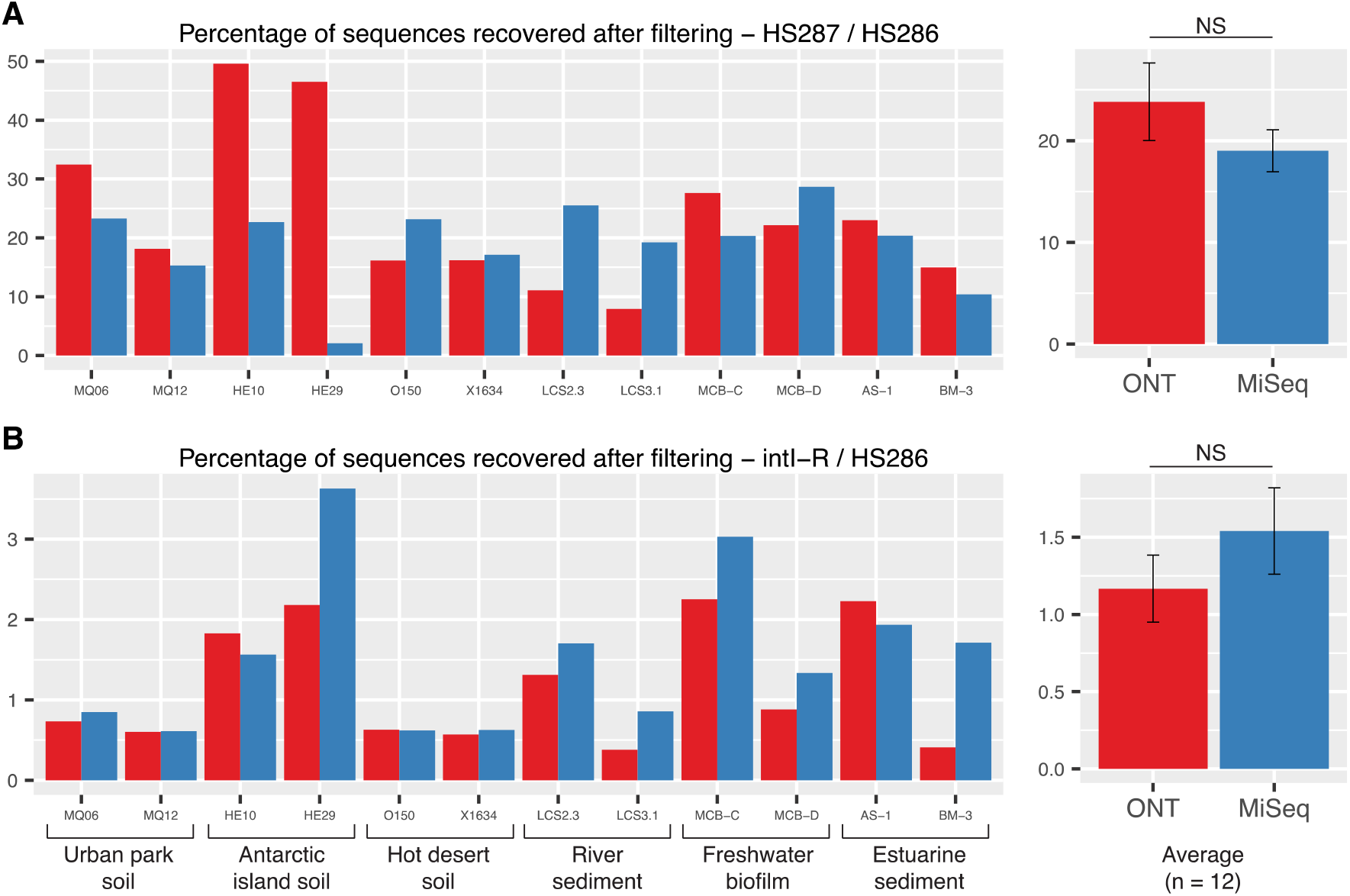
Percentages of sequences recovered after bioinformatic filtering. Filtering involved screening (**A**) HS287 / HS286 sequences for cassette recombination sites (*attC*s) and (**B**) intI-R / HS286 sequences for integron integrase (IntI) encoding genes to ensure that they represented amplicons of genuine integrons. Average (± 1 S.E) diversity for each analysis are shown on the right-hand side of each panel. Differences between Nanopore (ONT) and Illumina MiSeq technologies were not significant (NS) as determined by two-sample T-tests.

**Figure S2.**
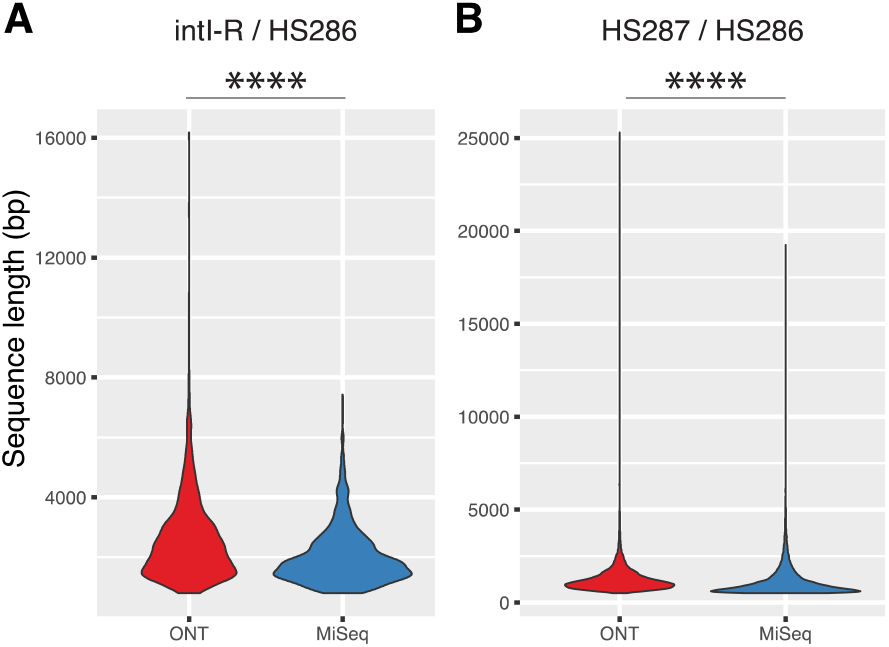
Sequence lengths recovered by each sequencing technology. The violin plots show the range of sequence lengths (bp) for (**A**) intI-R / HS286 and (**B**) HS287 / HS286 primer sets. The width of each curve represents the relative density of datum points across the ranges. For both primer sets, the average length of recovered amplicons (n=12) is significantly larger (Wilcoxon rank sum test, P<0.0001) when sequenced with Nanopore (ONT) compared MiSeq sequencing.

**Figure S3.**
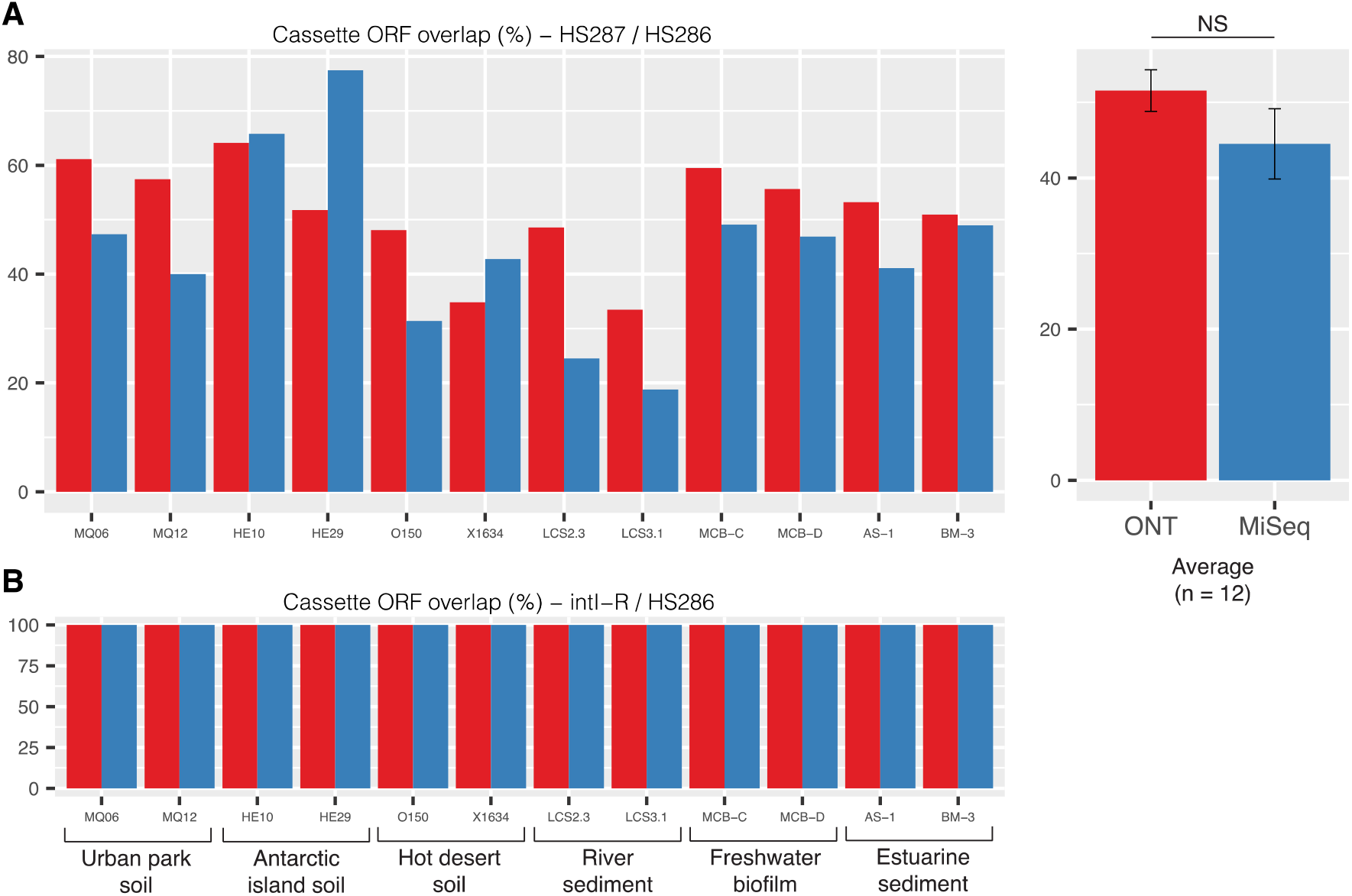
Overlap in recovered ORFs between Nanopore (ONT) and MiSeq sequencing technologies. The percentage of ORFs recovered from one sequencing technology that were covered by reads from the other technology are shown for (**A**) HS287 / HS286 and (**B**) intI-R / HS286 primer sets. ORFs considered to present in the opposite sequencing technology had to have a mean coverage depth of at least 1x that spanned at least 98% of the ORF. The average (± 1 S.E) percentage overlap for HS287 / HS286 data is shown on the right-hand side of panel (**A**). There was no significant (NS) difference between ONT and MiSeq (Two-sample T-test, P=0.209).

**Figure S4.**
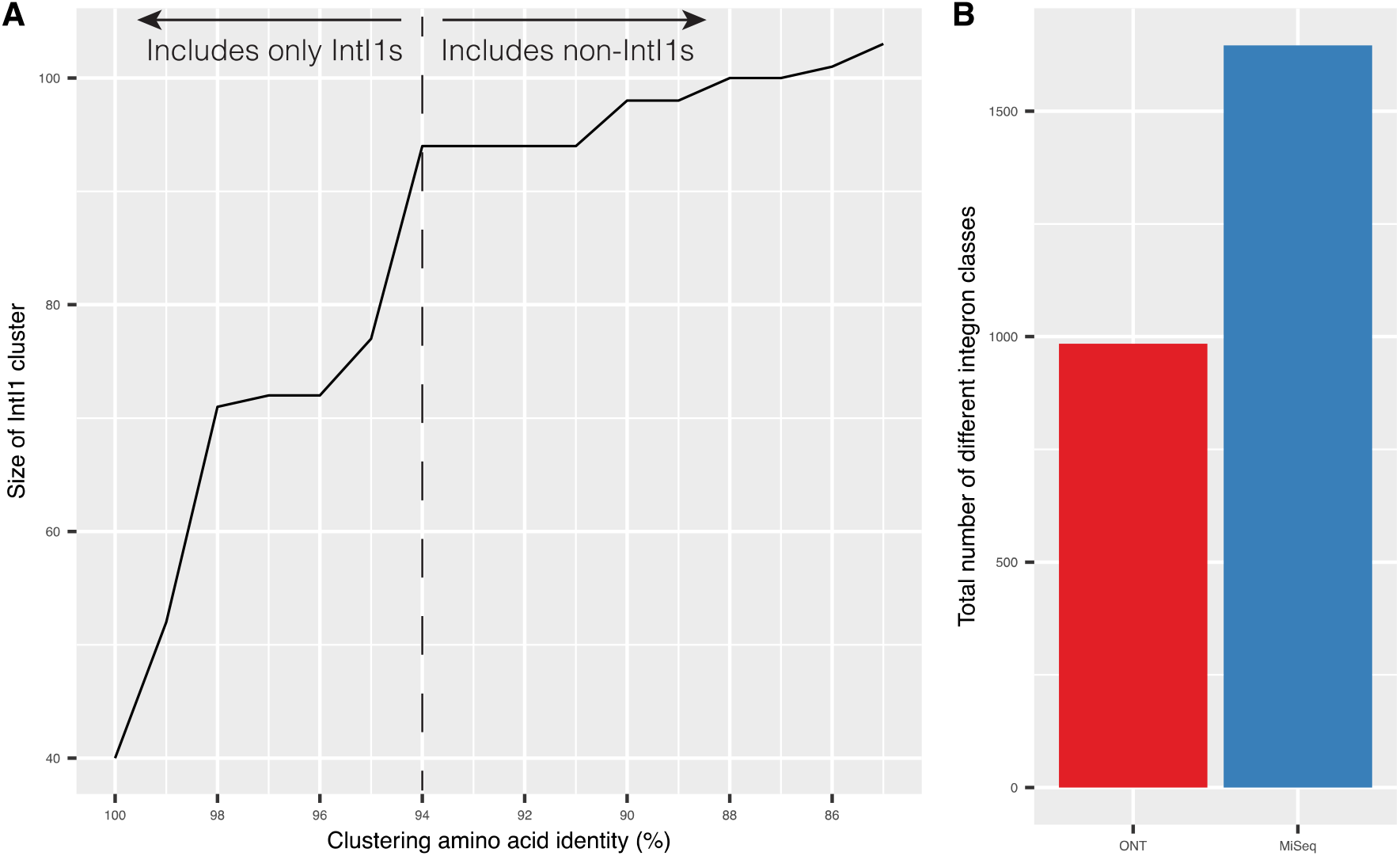
Diversity of integron classes. (**A**) The amino acid clustering threshold for integron classes was determined using class 1 integron integrases (IntI1) present in our dataset. Decreasing amino acid clustering thresholds were iteratively set until all IntI1s were grouped in the same cluster and all non-IntI1s were excluded. A protein sequence was considered to be IntI1 if it aligned with any previously characterised class 1 integron in GenBank using BLASTP (>98% amino acid identity and >70% subject cover). An amino acid clustering threshold of 94% was found to include all IntI1s (n=94) and exclude all non-IntI1s. (**B**) The total number of integron classes (based on a 94% amino acid clustering threshold) recovered for all samples (n=12).

**Figure S5.**
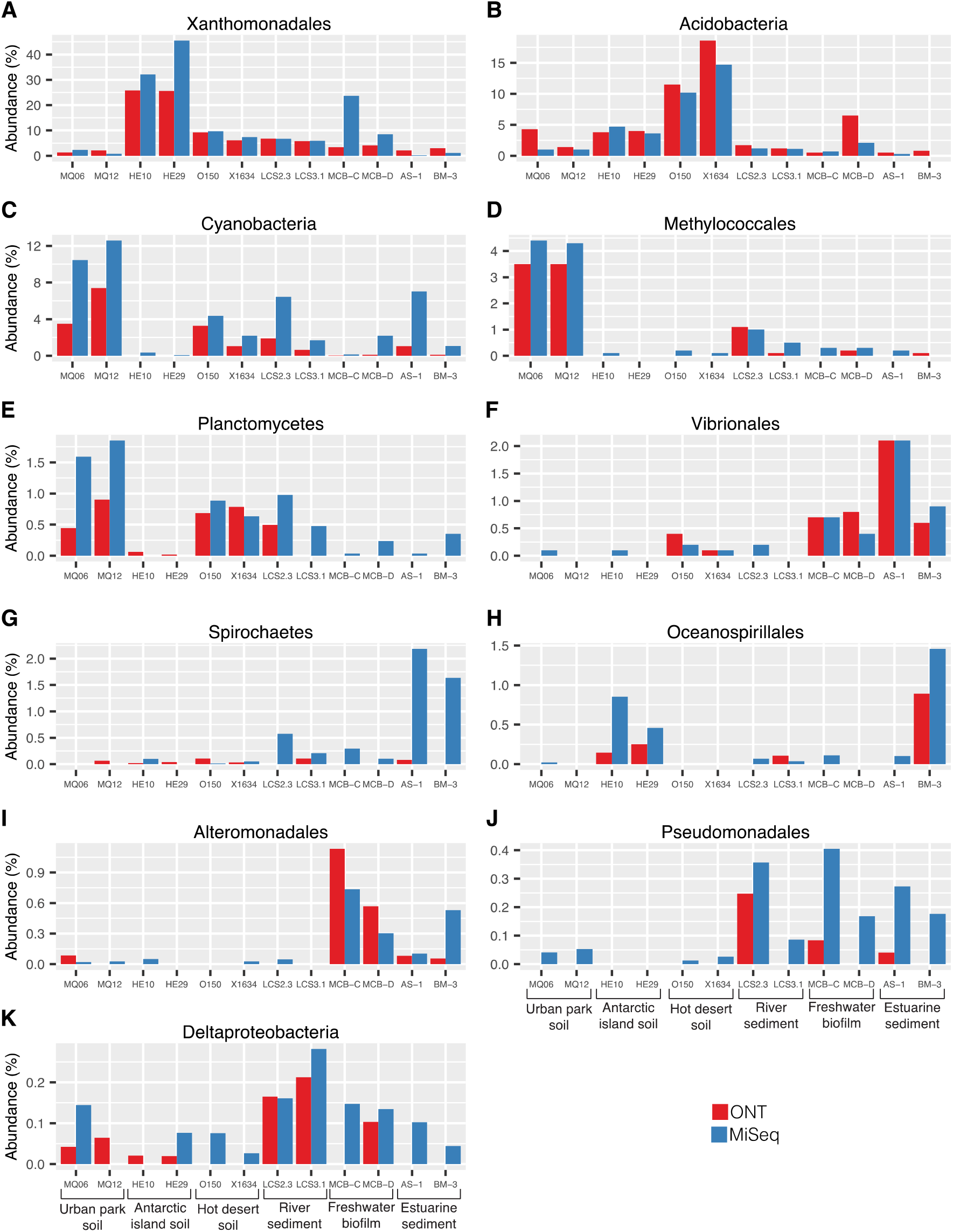
Taxonomic classification of gene cassette recombination sites (*attC*s). Taxonomic predictions are based on all eleven (**A-K**) available taxonomic models of chromosomal *attC*s. Each figure panel shows the proportion of *attC*s across each sample that exhibit sequence and structure conserved among that taxon.

